# Heat Conduction Simulation of Chondrocyte-Embedded Agarose Gels Suggests Negligible Impact of Viscoelastic Dissipation on Temperature Change

**DOI:** 10.1101/2024.02.08.579524

**Authors:** Erik Myers, Molly Piazza, Mark Owkes, Ronald K. June

## Abstract

Agarose is commonly used for 3D cell culture and to mimic the stiffness of the pericellular matrix of articular chondrocytes. Although it is known that both temperature and mechanical stimulation affect the metabolism of chondrocytes, little is known about the thermal properties of agarose hydrogels. Thermal properties of agarose are needed to analyze potential heat production by chondrocytes induced by various experimental stimuli (carbon source, cyclical compression, etc). Utilizing ASTM C177, a custom-built thermal conductivity measuring device was constructed and used to calculate the thermal conductivity of 4.5% low gelling temperature agarose hydrogels. Additionally, Differential Scanning Calorimetry was used to calculate the specific heat capacity of the agarose hydrogels. Testing of chondrocyte-embedded agarose hydrogels commonly occurs in Phosphate-Buffered Saline (PBS), and thermal analysis requires the free convection coefficient of PBS. This was calculated using a 2D heat conduction simulation within MATLAB in tandem with experimental data collected for known boundary and initial conditions. The specific heat capacity and thermal conductivity of 4.5% agarose hydrogels was calculated to be 2.85 J/g°C and 0.121 W/mK, respectively. The free convection coefficient of PBS was calculated to be 1000.1 W/m^2^K. The values of specific heat capacity and thermal conductivity for agarose are similar to the reported values for articular cartilage, which are 3.20 J/g°C and 0.21 W/mK (Moghadam, et al. 2014). This suggests that in addition to 4.5% agarose hydrogels mimicking the physiological stiffness of the cartilage PCM, they can also mimic the thermal properties of articular cartilage for *in vitro* studies.

## A. Introduction

### A1. Chondrocyte Homeostasis Depends on Loading and Temperature

Chondrocyte metabolism is linked to both physical loading (Lee, et al. 2018) such as exercise, as well as changes in whole-joint temperature (Newman, Bowles and Buckle 2017). Chondrocytes are encapsulated within their peri-cellular matrix within the extra-cellular matrix of cartilage, both of which are viscoelastic and dissipate some loading energy as heat during physical activity. Changes in environmental temperature affect the rates of enzymatic production of amino acid precursors and other important metabolites by decreasing the required activation energy for various central metabolism reactions. These metabolic products are used by chondrocytes to produce and maintain the extra- and peri-cellular matrices for cartilage homeostasis. Studies of chondrocyte mechanotransduction often encapsulate chondrocytes in viscoelastic hydrogels like agarose. However, during cyclical mechanical stimulation of encapsulate chondrocytes, it is unknown if heat generation is due exclusively to metabolic activity or viscous heat created through cyclical loading, and how similar this heat generation is to actual cartilage.

### A2. Cartilage and Agarose as Viscoelastic Materials

Cartilage is a viscoelastic material (Hayes and Bodine 1978). Cyclical loading during activities such as exercise will cause the cartilage to store and return some energy through its elastic response and dissipate the remainder of the energy as heat through its viscous response. Healthy human chondrocytes are surrounded by an ECM which has a complex modulus of around 1.5 MPa (Normand, et al. 2000). This release of heat will naturally cause a temperature change in the cartilage, and cartilage temperature increases by as much as 5-6°C in cases of rigorous exercise (Becher, et al. 2008). However, it is unknown how much of this increase is associated with viscoelastic dissipation and how much is associated with altered chondrocyte metabolism. Agarose hydrogels make a suitable *in vitro* model for chondrocyte mechanotransduction because at 4.5% weight per volume their complex modulus is similar (∼148 KPa) to the cartilage PCM (Jutila 2013) (Darling, et al. 2010). However, for chondrocyte mechanotransduction studies the thermal properties of agarose need to be characterized so that studies can distinguish between viscoelastic and metabolic temperature increases.

### A3. Heat Generated by Loading and Chondrocyte Metabolism

Prior studies show that agarose-embedded chondrocytes alter their central metabolism in response to applied loading (Zignego, Hilmer and June 2015). Thus, quantification of temperature during cyclical loading of agarose-embedded chondrocytes may reflect loading-induced changes in chondrocyte metabolism. However, to interpret temperature changes of agarose-embedded chondrocytes subjected to cyclical loading, the thermal properties of agarose are needed. Once these properties are known, then the observed temperature increases can be separated between those due to cyclical loading of the viscoelastic agarose and those due to chondrocyte metabolism. Thus, the goal of this study was to lay the foundation for the relationship between the mechanics of cartilage loading and changes to chondrocyte metabolism. To this end, a mathematical model was created that accurately simulates 2D heat diffusion through a representative agarose hydrogel and accounts for the temperature increase in the gel solely from viscoelastic loading.

## B. Methodology

### B1. 2D Heat Conduction in a Cylindrical Hydrogel

Heat diffusion through a 2D cylindrical viscoelastic solid is modeled using the heat conduction equation in cylindrical coordinates:

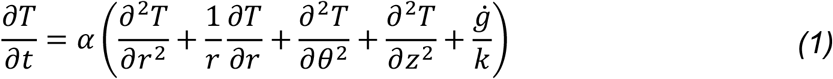

Above α is the thermal diffusivity of the solid and *g* is the heat generated in the material from viscoelastic loading, k is the thermal conductivity, and T is temperature. In the equation r, θ and z are the cylindrical coordinate directions. Because the boundary and initial conditions are uniform in the azimuthal direction, the equation is reduced to:

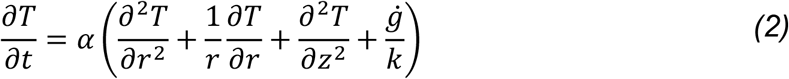

The convection coefficient and thermal diffusivity of the system are necessary to solve these equations. The thermal diffusivity is defined as:

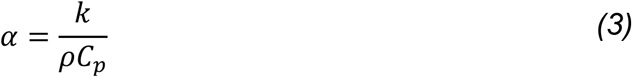

Where k is the thermal conductivity, *ρ* is the density and *C*_*p*_ is the heat capacity.

The model is validated by testing with conditions similar to tissue culture. Hydrogels are immersed in PBS at 37°C for a temperature boundary condition along the outside radius and bottom edge, as well as free air convection at 37°C on the top face. To solve computationally, Equation 2 was discretized (Figure 1, Equation 3):

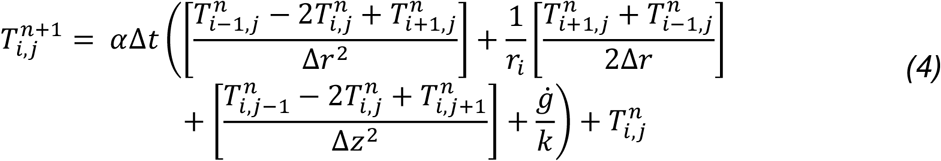

**Figure 1:**
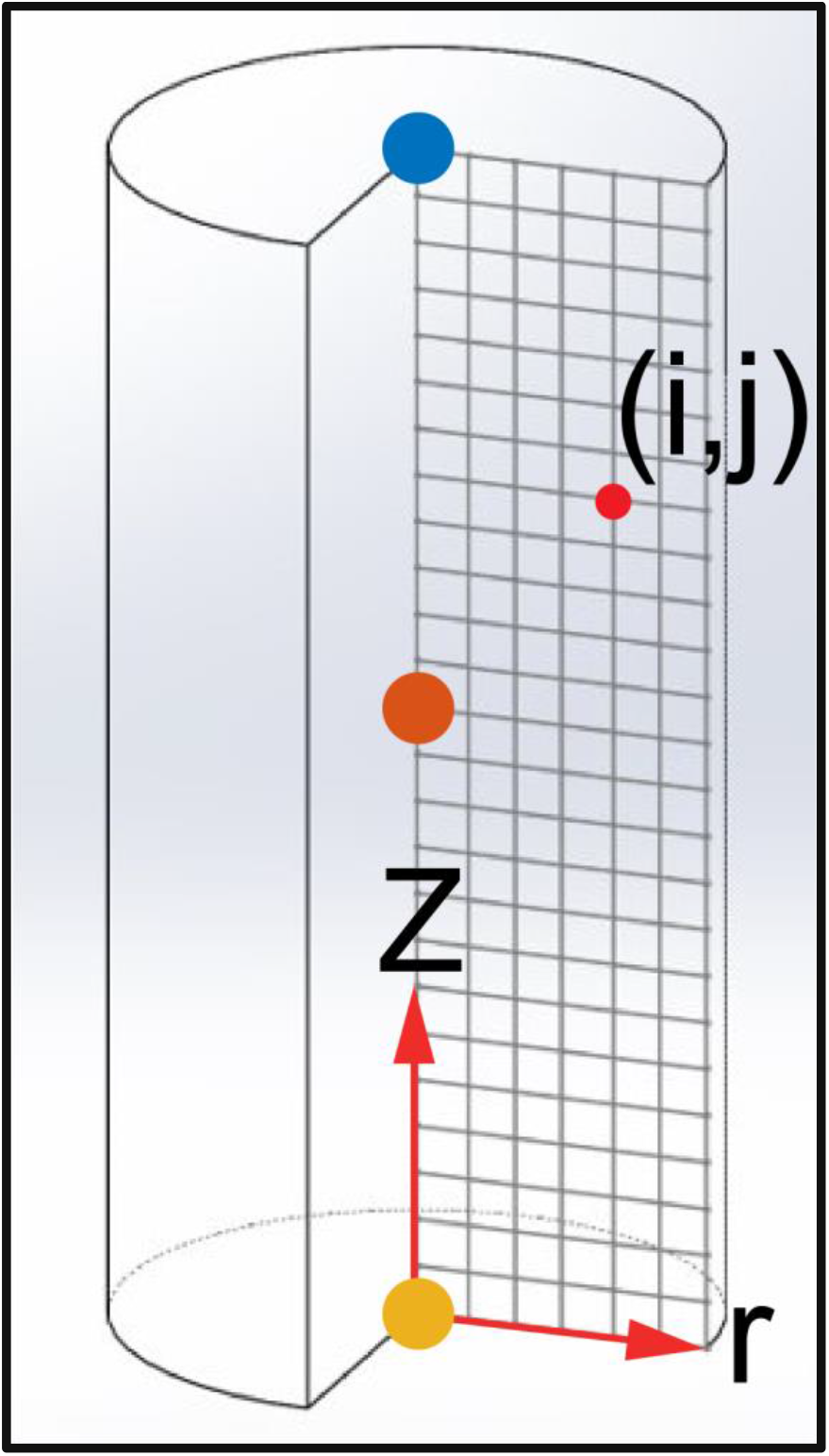
An example hydrogel cylinder is discretized using a uniform 2-D grid for simulation of heat diffusion. Since the boundary and initial conditions are symmetric about the azimuth direction a 2-D grid in r and Z is used. A uniform grid spacing is applied to simplify the discretization. Colored points represent locations of interest, including the top face (Blue), the midpoint (Orange), bottom face (Yellow), and current position (Red).

**Figure 2:**
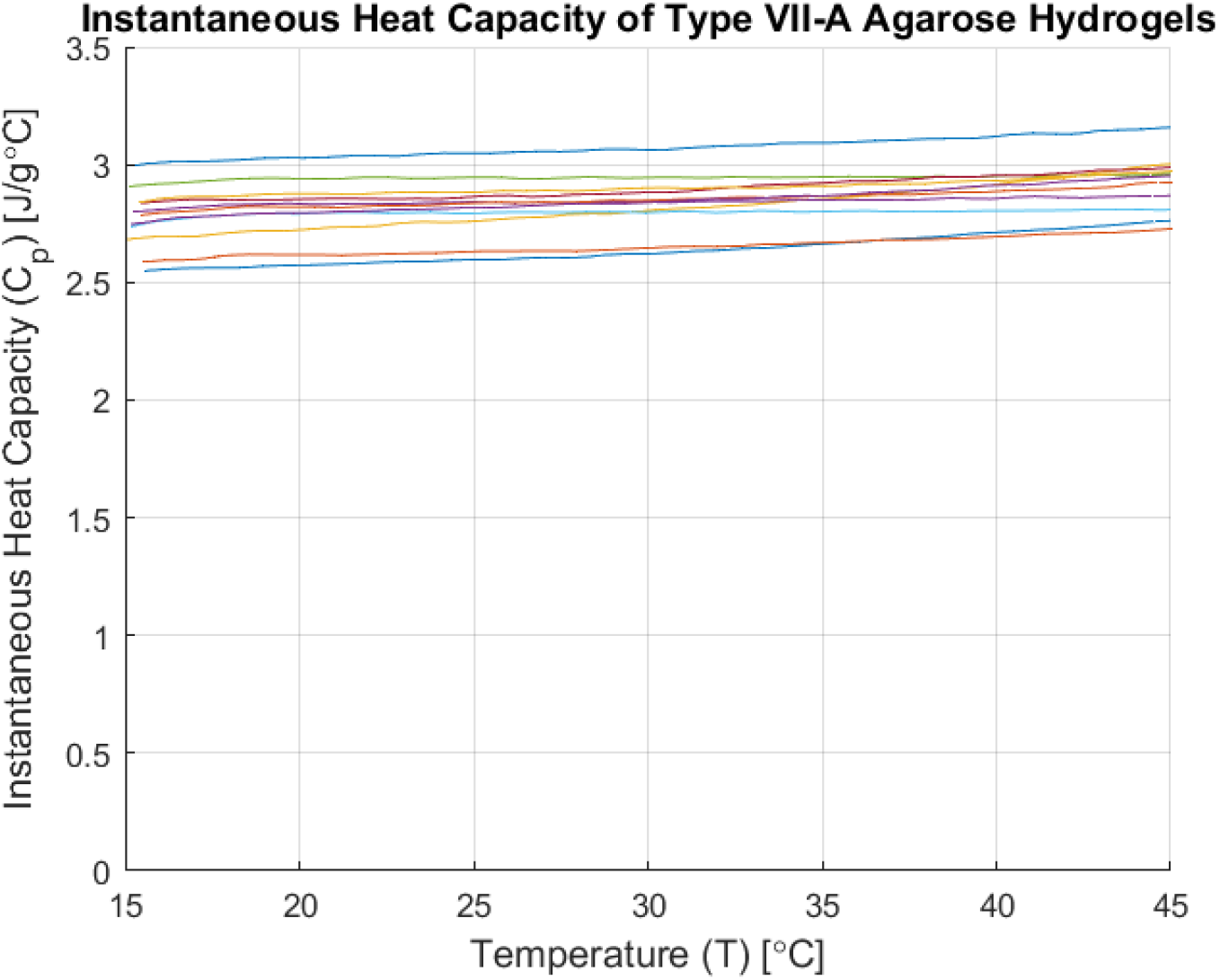
Instantaneous Heat Capacity of Agarose samples Across 15-45°C Calculated using DSC and a Constant Heat Ramp shows an average heat capacity of 2.85J/g°C. Each trace represents a single sample of 4.5% agarose gel from n=10 overall. DSC sample pans were filled with 30-45mg of agarose and sealed with cyanoacrylate. The DSC heated the sample pans and a reference pan from 15-45°C at a rate of 2°C/min while measuring the heat flow into both pans. This was used to calculate the instantaneous heat capacity at each temperature which is shown in this graph.

Above, *i, j* refers to the r and z coordinates of each point (i = 1,2,…,N_r_ and j = 1,2,…,N_z_), respectively, Δt is the time step, *n* is the current time point, Δr is the radial spacing of each node, and delta z is the height spacing of each node. The convection boundary conditions can be discretized to:

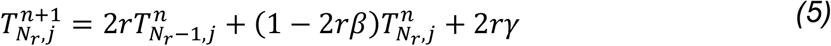

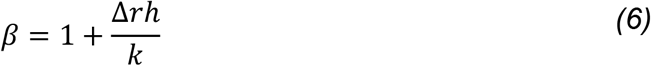

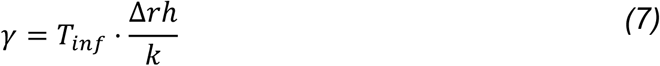

Above, *N*_*r*_ is the outermost point on the cylinder’s surface and *h* is the convection coefficient. These discretized equations can be iteratively solved in MATLAB to find the temperature distribution throughout the hydrogel at any time step. To determine the appropriate timestep, the following equation was used (Ozisik 1993):

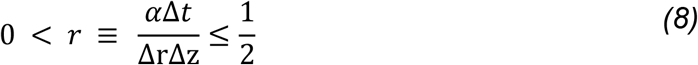

Which bounds the timestep between zero and the upper limit of stability (Ozisik 1993).

To complete the heat conduction simulation, several unknown parameters are needed. For agarose, the thermal conductivity and specific heat capacity were not known. In addition, the convection coefficient of air in the test chamber and the PBS that surrounds the hydrogels during testing were needed. These values were determined experimentally as follows.

### B2. Determination of Specific Heat Capacity

Samples of 4.5% type VII-a Agarose were added to aluminum sample pans and weighed (n=10). A hermetic seal was formed by applying Cyanoacrylate in equal amounts to both the sample pan and reference pan. Pan mass and adhesive mass were measured to be relatively constant between samples (Supplemental Table 1). Samples and reference pans were analyzed using Differential Scanning Calorimetry (DSC TA Instruments DSC2920) with a heating ramp from 15°C to 45°C at 2°C/min. Calibration was performed using a sapphire reference standard. The results of these DSC runs were further compared to an aluminum oxide standard to determine the specific heat of the agarose (Supplemental Figure 3). The outputs of the DSC were values of heat flow (*q*), temperature and time. This was converted into Heat Capacity (Equation 9):

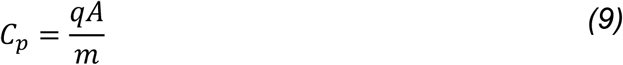

Above *m* is the mass of the sample in grams and *A* is a calibration coefficient specific to each machine that was found by running aluminum oxide under the same experimental settings as agarose.

### B3. Thermal Conductivity Testing

4.5% Low-gelling temperature agarose samples 0.25 inches thick and 2.5 inches in diameter were tested in a thermal conductivity machine following ASTM standard C177. To validate the thermal conductivity machine and ensure proper calibration of the thermistors and heating system, samples of neoprene foam insulation with a known thermal conductivity were used to create a profile of *k vs T* values. A plot of these values is available in Supplemental Figure 4. To test the agarose samples, the cold plate was held at 17°C while the hot plate ranged from ∼33-40°C. Power delivered to the hot plate was recorded and used to calculate heat flux through the agarose puck (Equation 10). This flux was then used to calculate the thermal conductivity of the material over the given temperature range.

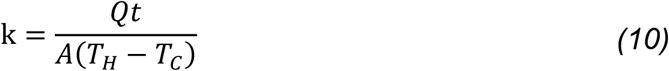

**Figure 3:**
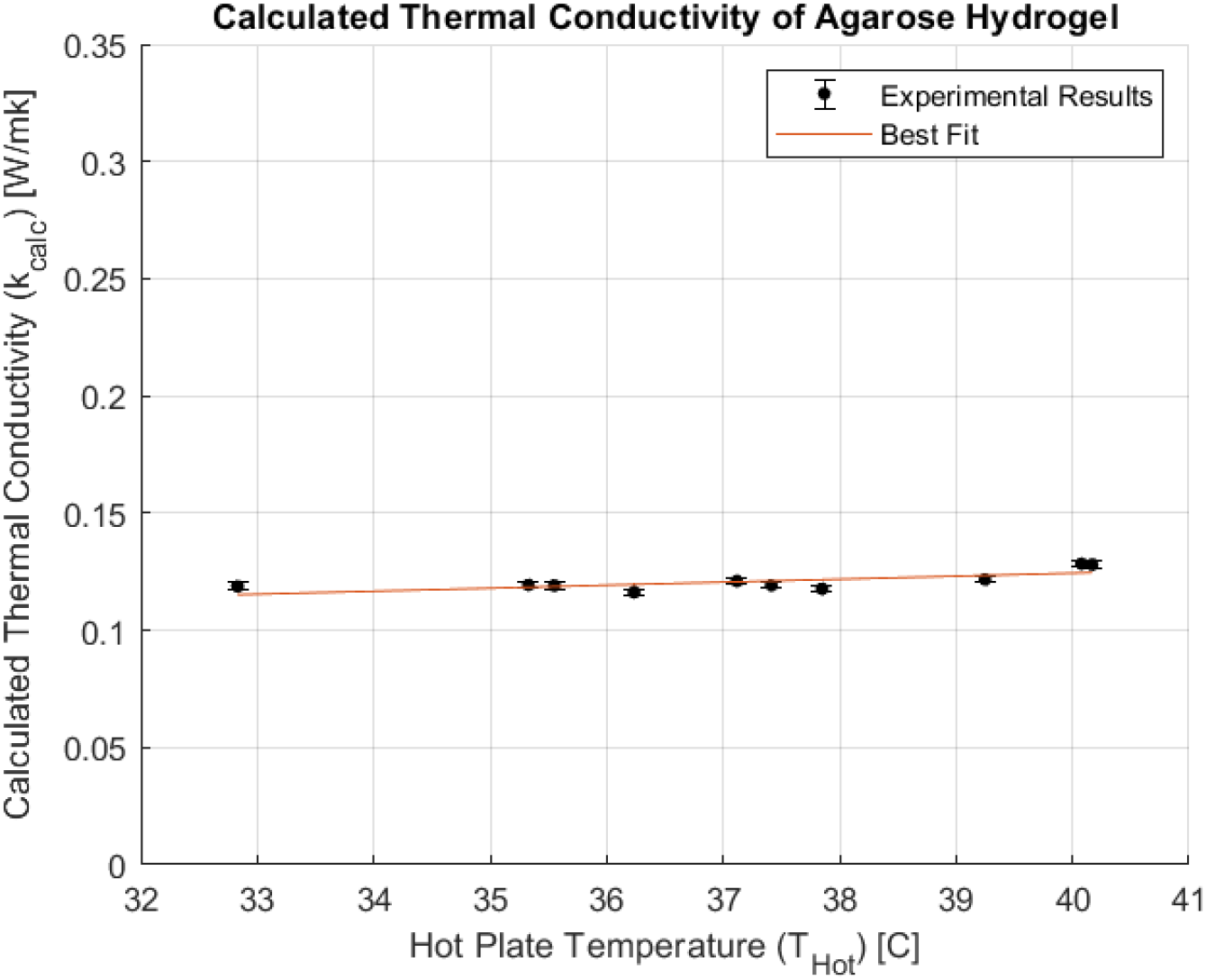
Thermal conductivity testing of 4.5% agarose hydrogel pucks following ASTM C177 shows a steady value of 0.121 w/mk across the temperature range of interest including 37°C. Each point represents a single test performed at a different hot plate temperature, with the error bar representing the standard error across the sample group. All ten tests were performed on the same agarose hydrogel puck.

**Figure 4:**
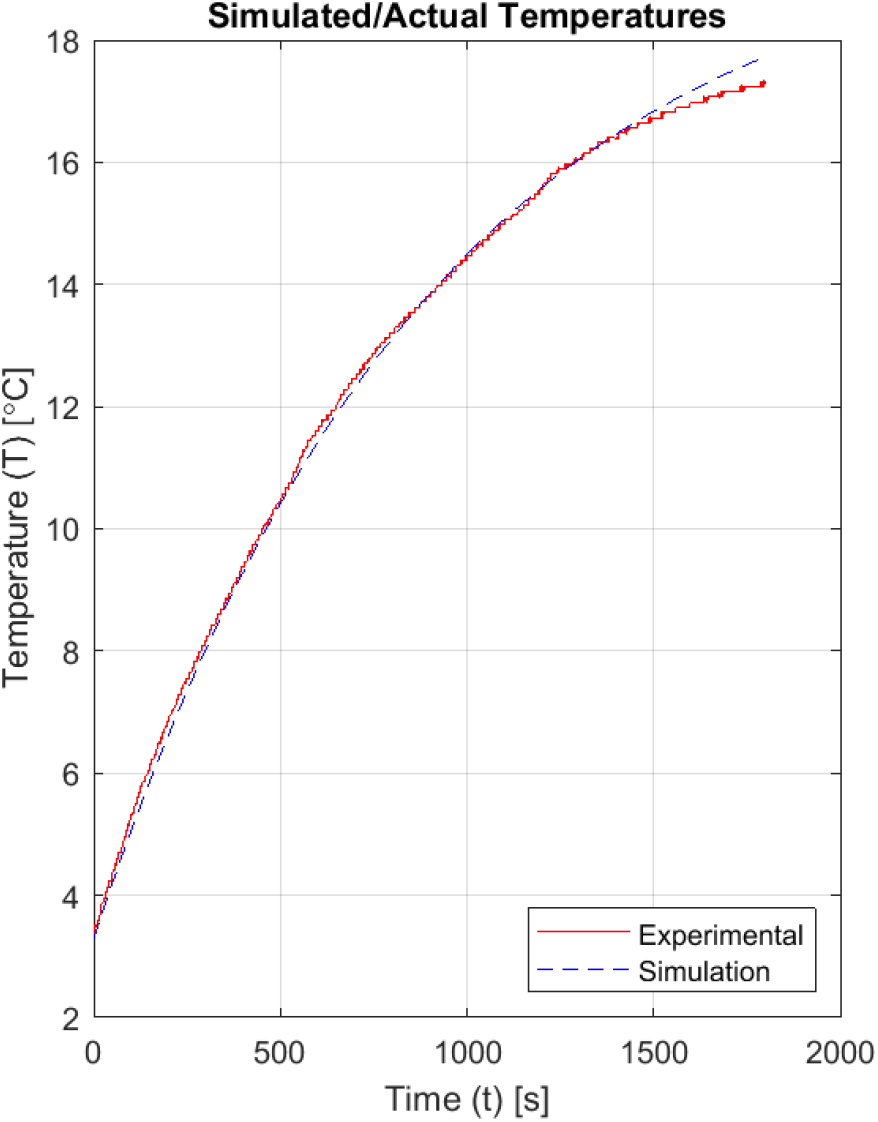
Simulated and experimental temperature vs. time using determined convection coefficient of air. The convection coefficient of air was determined by fitting the simulation to experimental data through iteratively adjusting the convection coefficient value until the error between the simulated and experimentally measured temperature converged.

**Figure 5:**
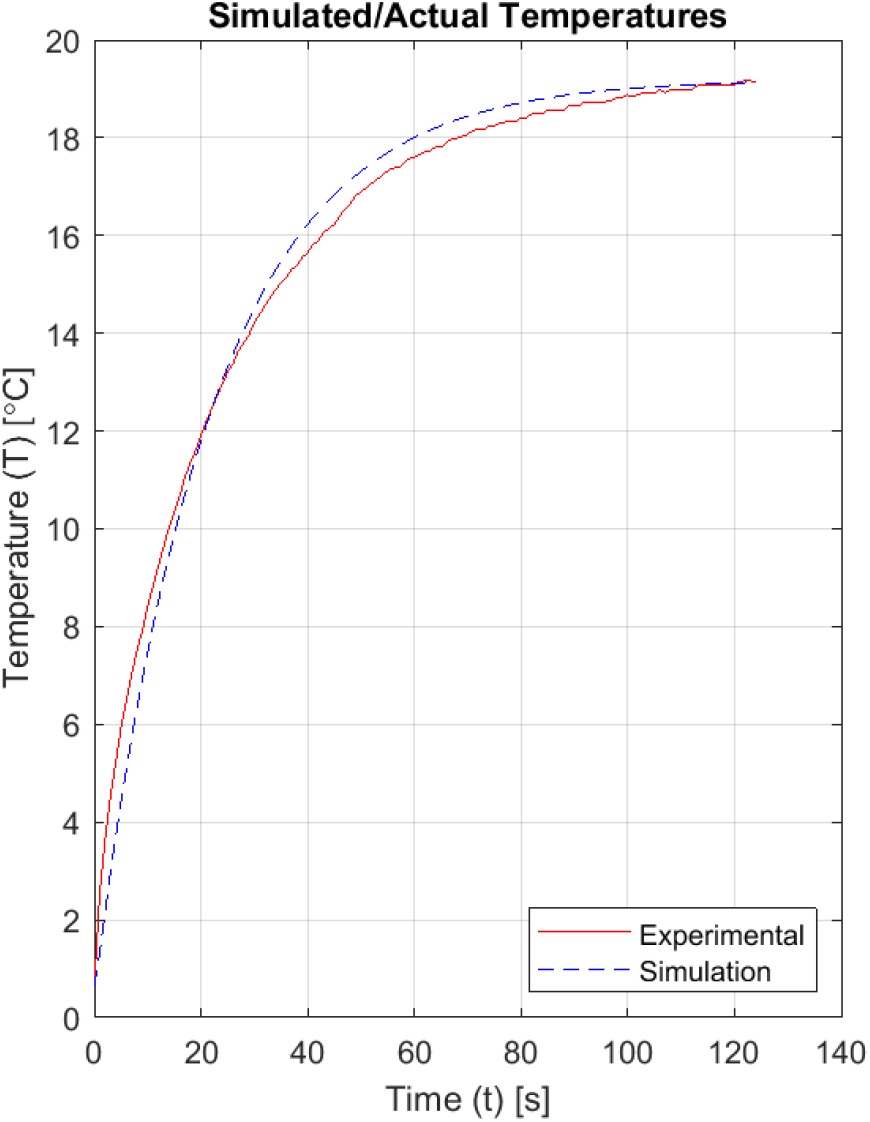
Calculation of the convection coefficient of PBS. The PBS convection coefficient was calculated as in Figure 4 using the now-determined h_air_ and then fitting the model to the experimental data by minimizing the SS_error_ between simulated and experimental temperature data. Results show the simulation using the final value of h_PBS_ and the experimental data.

### B4. Measurement of Convection Coefficients of Air and PBS

Convection coefficients for air and PBS were determined using a steel rod ∼2in long and ∼0.5in in diameter with known thermal characteristics (k = 51.9W/mK, C_p_ = 486 J/kgK, ρ = 7870 kg/m^3^). To measure the temperature at the center of the rod, a thermistor was embedded in a pre-drilled hole and sealed with thermal paste (AOS Thermal Compounds, Eatontown, NJ). To calculate the convection coefficient of air, the rod was cooled to 3°C then hung in stagnant room temperature air until the center temperature reached a steady-state value (T_inf_ = 20°C). To calibrate the simulation, initial bounding values for the air h-value were assumed. Then a bisection method was used to minimize the sum of squares of the error (SS_error_) between the actual temperature plot of the center of the rod compared to the theoretical temperature given the correct h-value for air. Once the convection coefficient of air was calculated, the rod was cooled to 1°C and submersed in room temperature PBS with the top face exposed to air for ∼140s. This follow up experiment was performed knowing all the thermal characteristics of the steel as well as the convection coefficient of air. The data from this experiment was used to find the only remaining unknown: the convection coefficient of PBS. As above, a bisection method was used to narrow the acceptable h-value of PBS until the error between the experimental and theoretical plots was minimized.

### B5. Simulated Viscous Heat Generation

The viscoelastic component of heat generation was added to the simulation using the equation (Esmaeelia, et al. 2018):

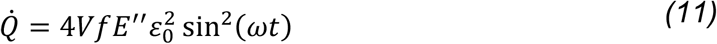

Where *V* is the volume of the agarose, *f* is the frequency of loading, E’’ is the loss moduli, *ε*_*0*_ is the maximum amplitude of strain, *ω* is the frequency in rad/sec, and *t* is time.

Once *k*_*agarose*_, *C*_*p_agarose*_, *h*_*air*_, and *h*_*PBS*_ were determined, the MATLAB iterative solver simulated the thermal behavior of an agarose hydrogel at various initial temperatures, environmental temperatures, and boundary conditions. To mimic the environmental conditions in a tri-gas incubator, the top boundary condition was set to free air convection, and the remaining boundaries were set to free PBS convection, both at *T*_*inf*_ = 37°C. The appropriate upper timestep limit was calculated to be *Δt* = 0.06s. The initial hydrogel temperature was set to 37°C, and the agarose temperature distribution as well as material properties were assumed to be isotropic. This setup allowed for an accurate prediction of the effect of the viscoelastic loading response on the overall temperature of the hydrogel.

## C. Results

### C1. DSC Calculation of Specific Heat Capacity

To determine the specific heat capacity of agarose, the heat capacity data were plotted with respect to temperature (Figure 2).

The calculated heat capacity of agarose over the entire range of temperatures was 2.85J/g°C with a standard error of 0.04J/g °C. These data represent the average heat capacity of each of the n=10 samples over the temperature range, and then taking the average of those 10 values.

### C2. Thermal Conductivity Calculation

The data from the thermal conductivity tests were plotted with respect to the hot plate temperature (Figure 3). The calculated thermal conductivity of agarose at 37°C is 0.121W/mK.

This plot yields temperature-dependent results for the agarose over a range of values which capture average joint temperature. Linear interpolation yields the following equation:

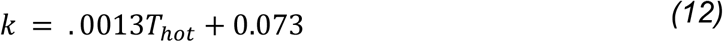

With an average thermal conductivity over the entire range of 0.121±0.001 W/mK. Note that this value is different from reported thermal conductivity values of cartilage (0.21 W/mK) (Moghadam, et al. 2014).

### C3. Convection Coefficient Calculations

The simulation result using the final calculated value for the convective coefficient h_air_ fits the experimental results well (Figure 4). The final calculated h_air_ value was h_air_ = 23.2 W/m^2^K. The full table of calculated values and errors can be found in the supplemental material.

For the PBS convection coefficient, h_PBS_ was similarly determined by reducing SS_error_ and plotting the results. The final calculated value was h_PBS_ = 1000.1 W/m^2K.

### C4. Viscoelastic Heat Generation

After running the MATLAB solver, temperatures at specific locations on the hydrogel were plotted to analyze the effects of viscoelastic heating during cyclical loading:

Three separate temperatures are plotted: along the centerline of the hydrogel, at the bottom face along the PBS boundary condition, and along the top face at the air boundary condition (Figure 6). As expected, temperature at the PBS boundary is much less susceptible to viscoelastic effects than the air or interior points. The overall temperature increase along any of these profiles was negligible during sinusoidal compression. Zooming in on the temperature profile of the bottom face (Along the PBS boundary) reveals the more apparent effect of sinusoidal viscous heat generation (Figure 7).

**Figure 6:**
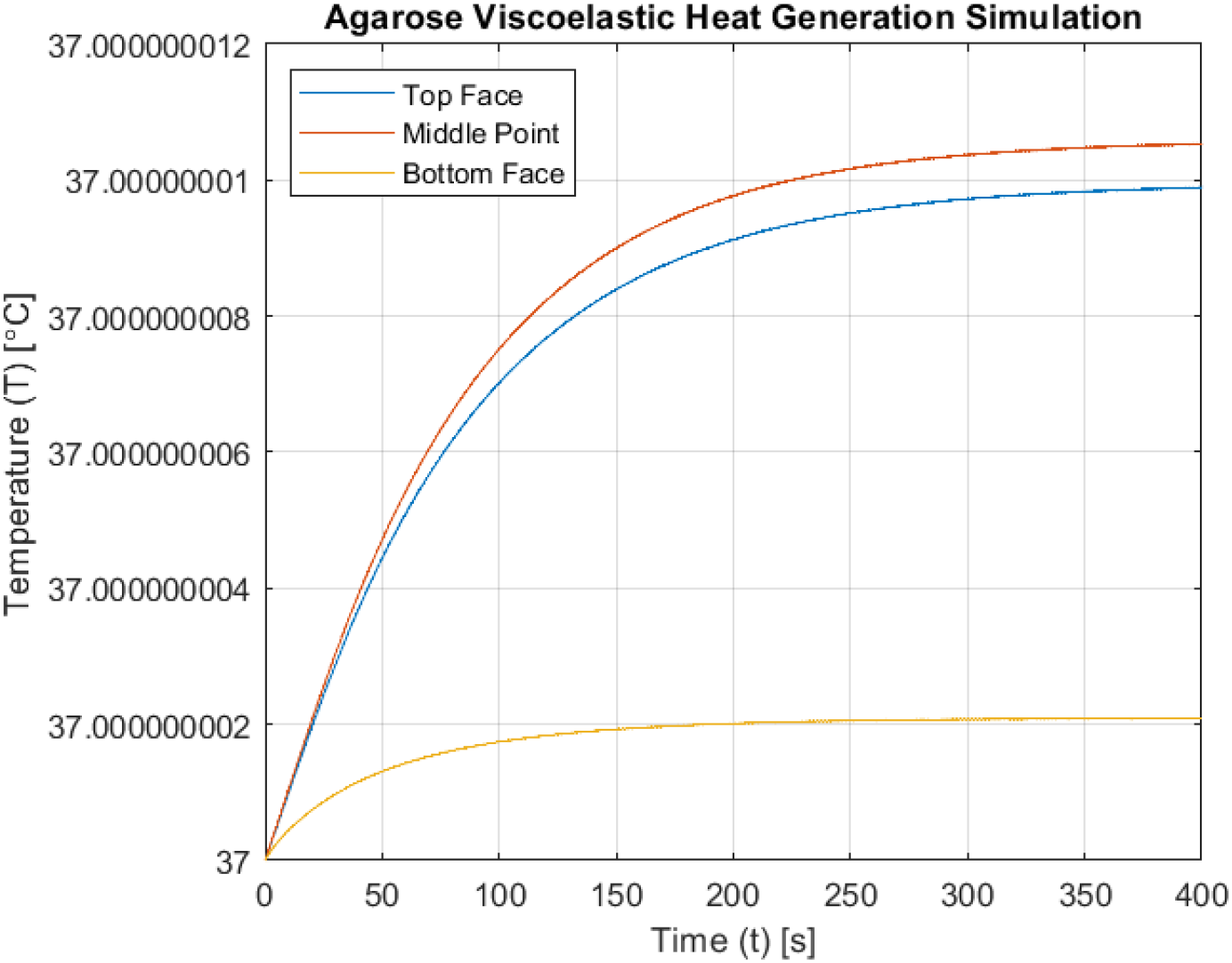
Simulated viscous heat generation for steady state agarose at 37°C with 5% compressive strain sinusoidal loading at 1.1Hz. Temperature changes at the center of the hydrogel were as little as 1e-8°C. The temperature effect of heat generation is different at the surfaces of the hydrogel due to the convective effect of the OBS surrounding the hydrogel. Since the temperatures of the PBS and air, T_inf_, are dictated within the code, the fluid temperature never changes, making them a heat sink during heat generation within the gel. Surfaces such as the top and bottom face which are in direct contact with the fluids therefore have a temperature closer to each fluid’s T_inf_ during loading.

**Figure 7:**
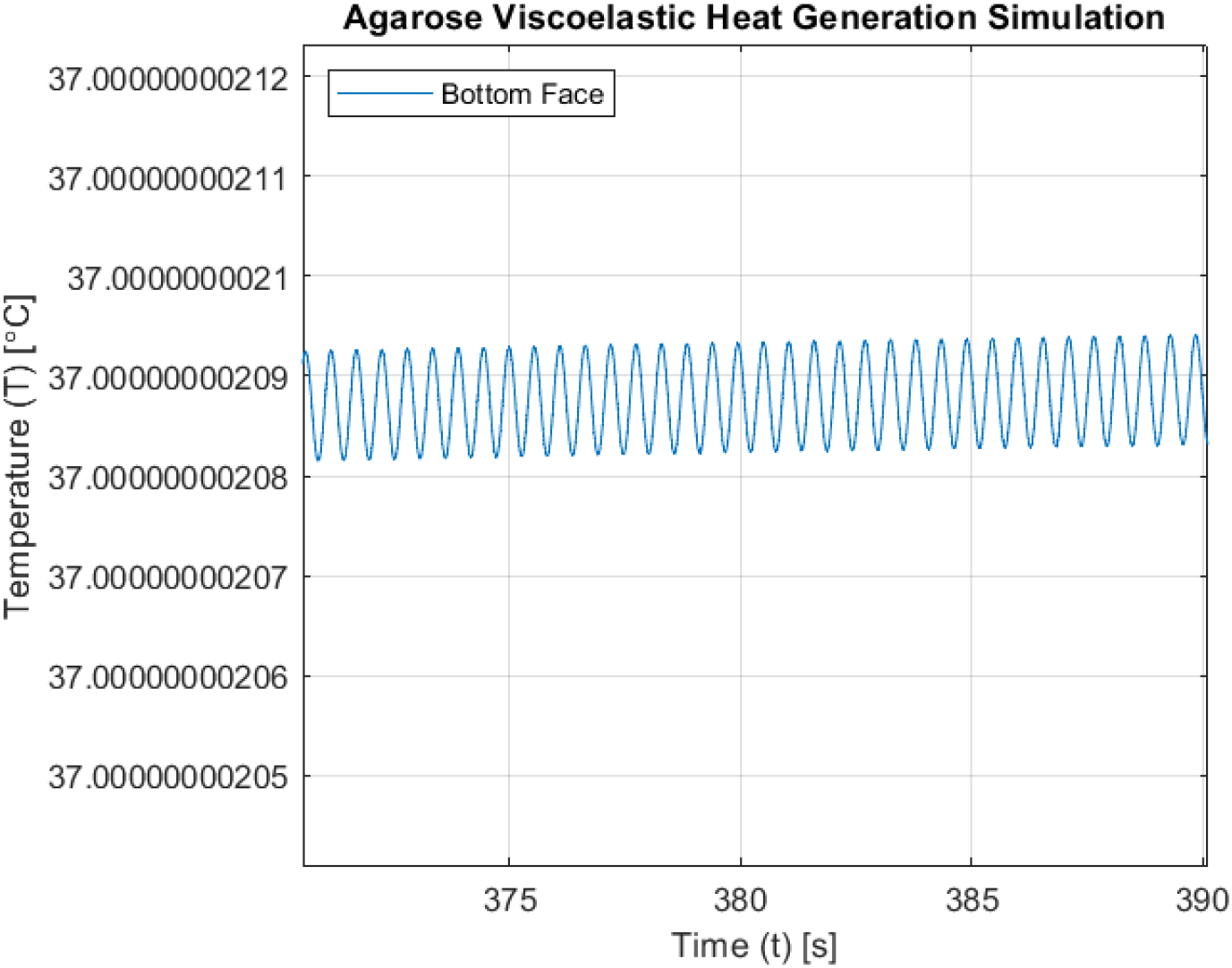
A high-resolution plot of simulated agarose temperature over time under cyclic loading conditions shows continued oscillations. The simulation started with both the agarose and PBS at a steady state 37°C and applied loading 5% compressive loading at 1.1Hz. These oscillations occur at twice the frequency of the loading that is applied. The temperature increases due to viscoelastic loading both as the agarose is compressed and as it dissipates viscous energy. The temperature decreases due to the PBS around the hydrogel acting as a heat sink.

While the actual temperature increase at the boundary is limited due to the convection boundary condition, the sinusoidal heat generation can be seen affecting the temperature for every loading cycle. The frequency of the heat generation is double that of the loading as energy is lost to heat dissipation during both the loading and unloading sections of the loading cycle.

## D. Discussion

### D1. Viscoelastic Loading has a Negligible Effect on Agarose Temperature

These data and simulations show that the low heat capacity of the agarose allows heat generation from cyclical loading to be diffused into the PBS rapidly. This simulated viscous heat generation reaches a steady state negligibly greater than the original temperature, and this increase (<10^−8^C) is undetectable by current instrumentation. The effect of viscoelastic dissipative heat generation on the agarose is most detectable at the center of the cylindrical samples. Future studies using cells for mechanotransduction experiments with these conditions are justified in neglecting agarose-based viscous heat generation as a source of substantial temperature change.

### D2. The Use of Agarose as an *in vitro* Cartilage Model

The thermal properties of 4.5% low-gelling temperature agarose differ from that of cartilage. Cartilage has a specific heat of 3.20J/g°C (Moghadam, et al. 2014) which is greater than the 2.85J/g°C of this agarose. This difference affects how potential heat generation from chondrocyte metabolism and viscoelastic loading is transported into the surrounding environment. Therefore, while measuring the temperature increase of agarose may be a useful tool for analyzing heat production by changes in chondrocyte metabolic activity, the results of such a measurement cannot be directly applied to cartilage. Creating a hydrogel with a heat capacity closer to that of healthy cartilage would allow a closer replication of *in vivo* chondrocyte habitat and a more accurate representation of thermal behavior during loading.

### D3. Implementing a 2D Heat Diffusion Model in Agarose to Determine Heat Generated by Chondrocytes

The present implementation of a 2D heat diffusion model allows for the dynamic comparison of temperature changes in an agarose hydrogel from several inputs including environmental factors, and viscous loading response (code in supplemental material). With these values accounted for any remaining heat generation can be assumed to be due to chondrocyte metabolism. Understanding heat transfer in cell-embedded agarose is important to analyze temperature changes that may result from cellular mechanotransduction. These data and model allow for analysis of potential interactions between macroscale temperature changes and metabolic reaction rates within chondrocyte central metabolism.

Viscous heat generation appears to have a negligible effect on the temperature of agarose during cyclical compression. This is because the hydrogels are submersed in PBS during compression. The PBS acts as an effective heat sink and maintains the agarose at a stable temperature during testing. To measure the temperature change of agarose during sinusoidal compression to assess chondrocyte metabolism, it may be necessary to embed a thermistor within the sample to minimize boundary effects on measured temperatures. Future studies can build on these results to assess sample temperature as a surrogate measure of metabolic activity during mechanotransduction studies.

## Funding Acknowledgement

This study was funded by the NIH (NIAMS R01AR073964 and R01AR081489) and NSF (CMMI 1554708).

