## Supplemental File 1 for "Heat Conduction Simulation of Chondrocyte-Embedded Agarose Gels Suggests Negligible Impact of Viscoelastic Dissipation on Temperature Change"

#### Supplemental Material

| Sample | Pan Mass (mg) | Sample Mass (mg) | Sealant Mass (mg) | Sealant Mass Variance (mg) | Average Cp (J/g℃) |
|---|---|---|---|---|---|
| 1 | 28.8 | 30.7 | 11.3 | 0.8 | 2.91 |
| 2 | 28.7 | 33.6 | 10 | 0.5 | 3.08 |
| 3 | 28.1 | 31 | 4.5 | 0.2 | 2.86 |
| 4 | 28.3 | 34.3 | 4 | 0.3 | 2.91 |
| 5 | 26.9 | 32.4 | 14.4 | 0.4 | 2.65 |
| 6 | 28.3 | 41.1 | 3.8 | 0.5 | 2.86 |
| 7 | 27.1 | 34.8 | 16.7 | 0.9 | 2.66 |
| 8 | 28.7 | 33.5 | 4.9 | 1.3 | 2.85 |
| 9 | 28.4 | 39.7 | 4 | 2.2 | 2.91 |
| 10 | 28.2 | 44.8 | 4.6 | 1.6 | 2.80 |
| Average ± SE | 28.2±0.2 | 35.6±1.5 | 7.8±1.5 | 0.9±0.2 | **2.85±0.04** |

*Supplemental Table 1: Mass Properties of Agarose DSC Pans and Measured Heat Capacity ($C_p$) of Agarose*

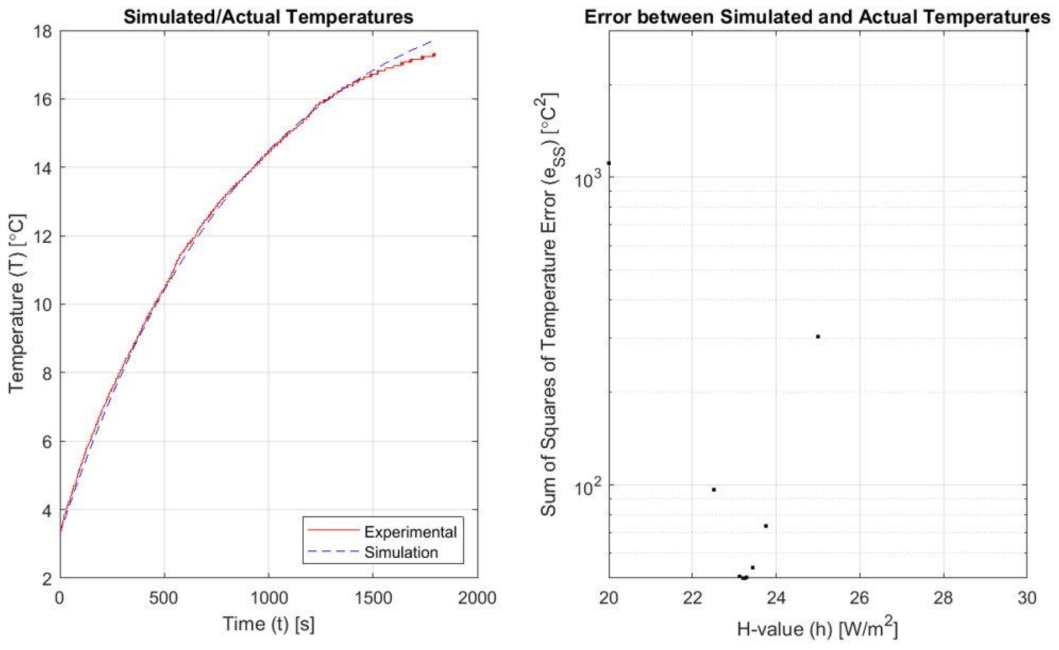

*Supplemental Figure 1: Simulated and Actual Cylinder Temperatures Vs Time (left) and Effect of Convection Coefficient of Air on Error Between Simulated and Actual Temperatures (right)*

| **h_value ($W/m^2$)** | **SS_Error (°$C^2$)** | **Calculation time (s)** |
|---|---|---|
| 20 | 1114 | 157 |
| 30 | 3003 | 121 |
| 25 | 304 | 141 |
| 22 | 97 | 130 |
| 24 | 74 | 132 |
| 23 | 51 | 121 |
| 23 | 54 | 132 |
| 23.28 | 50.03 | 131 |
| 23.20 | 49.74 | 131 |
| 23.24 | 49.75 | 140 |
| 23.22 | 49.71 | 131 |
| 23.21 | 49.72 | 158 |

*Supplemental Table 2: Minimization of Error by Iterative Calculation of Convection Coefficient of Air*

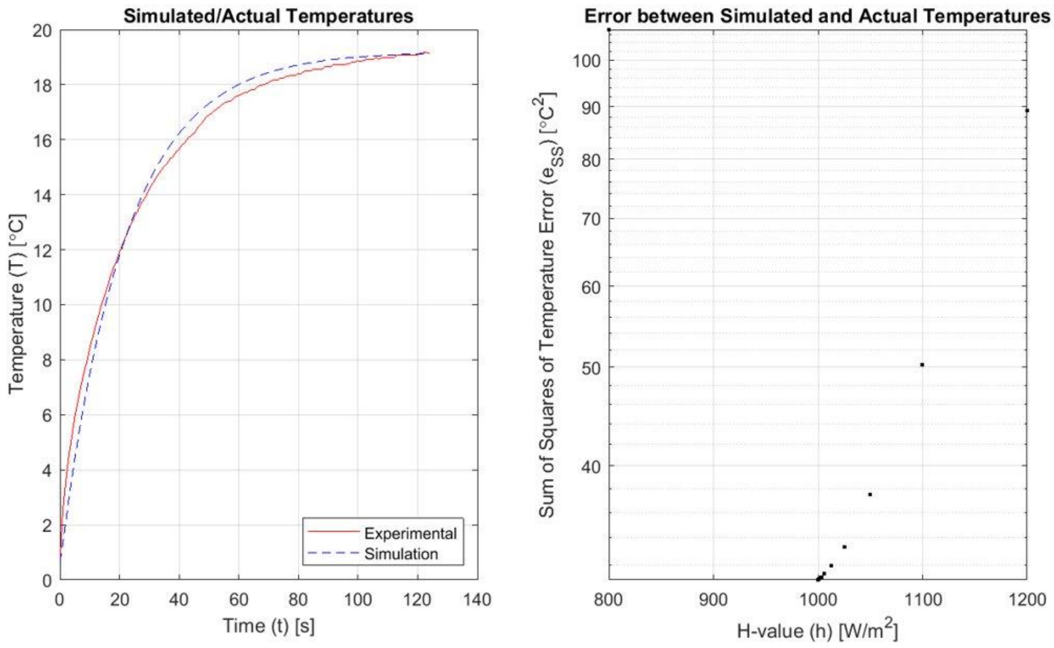


*Supplemental Figure 2: Simulated and Actual Cylinder Temperatures Vs Time (left) and Effect of Convection Coefficient of PBS on Error Between Simulated and Actual Temperatures (right)*

| h_value ($W/m^2$) | SS_Error ($°C^2$) | Calculation time (s) |
|---|---|---|
| 800 | 107 | 51 |
| 1200 | 89 | 40 |
| 1000 | 31 | 44 |
| 1100 | 50 | 43 |
| 1050 | 37 | 44 |
| 1025 | 33.35 | 44 |
| 1012.50 | 31.91 | 43 |
| 1006.25 | 31.36 | 43 |
| 1003.12 | 31.13 | 44 |
| 1001.56 | 31.03 | 44 |
| 1000.78 | 30.98 | 43 |
| 1000.39 | 30.95 | 43 |
| 1000.20 | 30.94 | 44 |
| 1000.10 | 30.94 | 43 |
| 1000.05 | 30.93 | 43 |

*Supplemental Table 2: Minimization of Error by Iterative Calculation of Convection Coefficient of PBS*

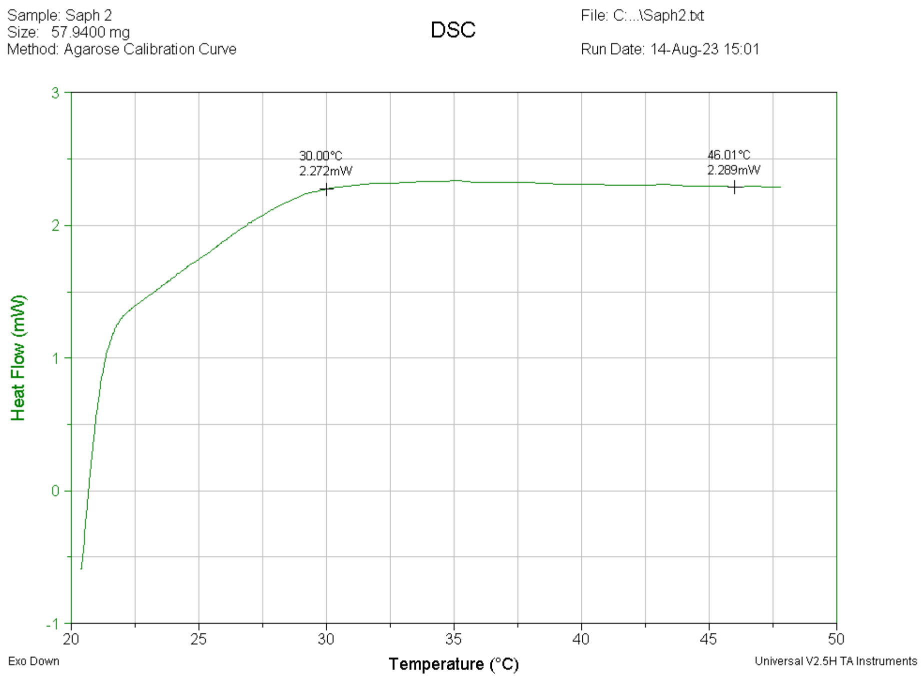


*Supplemental Figure 3: DSC Calibration with Aluminum Oxide*

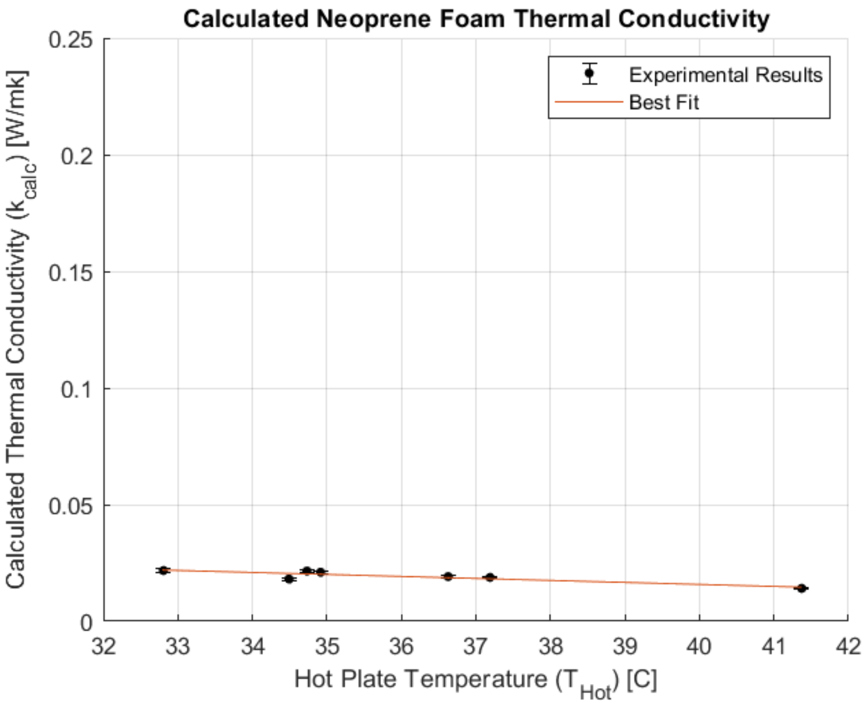


*Supplemental Figure 4: Thermal Conductivity Calibration with Neoprene Foam*

*Supplemental Code*

#### MATLAB 2D Heat Conduction Simulation Code

```
%% Erik Myers
% June Research Lab
% Montana State University

% 2D Transient Heat Conduction Simulation with Heat Generation
% V1.0 10/03/2023

% This code is the third version of the transient heat generation
% simulation, using Osizik's textbook and a modified setup of the
% two-dimensional steady-state heat conduction equation for cylindrical heat
% flow. This particular flavor has been modified to include heat generated
% from viscoelastic loading of the agarose gel.

%% Pre-Prep
clear
clc
close all
warning('off')

tic;    % Start timing the program for display later

%% User Defined Inputs
% This section is for the user to define anything they want about the
% simulation which may be changed frequently.

% Material Properties
    % Agarose Gel Material Properties
        Material = '4.5% Agarose Gel';

        switch Material
            case '4.5% Agarose Gel'
            R = (0.25/2) * 0.0254; % [m] Gel radius
            H = (0.5)    * 0.0254; % [m] Gel height
            k = 0.1;               % [W/mK] Thermal conductivity
            C_p = 2850;            % [J/kgK] Specific heat capacity, from the DSC calculations
            rho = 1069;            % [kg/m^3] Density
             alpha = k/(rho*C_p);   % [m^2/s] Thermal diffusivity

            case 'Aluminum'
            R = (0.562/2) * 0.0254; % [m] Cylinder radius
            H = (2)    * 0.0254;    % [m] Cylinder height
            k = 167;                % [W/mK] Thermal conductivity
            C_p = 896;              % [J/kgK] Specific heat capacity
            rho = 2710;             % [kg/m^3] Density
             alpha = k/(rho*C_p);   % [m^2/s] Thermal diffusivity

            case 'Steel'
            R = (0.511/2) * 0.0254; % [m] Cylinder radius
            H = (2)    * 0.0254;    % [m] Cylinder height
            k = 51.9;               % [W/mK] Thermal conductivity
            C_p = 486;              % [J/kgK] Specific heat capacity
            rho = 7870;             % [kg/m^3] Density
            alpha = k/(rho*C_p);    % [m^2/s] Thermal diffusivity

        end

        T_int = 22;           % Initial cylinder temperature

        % g = 0;              % Heat generation term. Commented for this code flavor

    % PBS Material Properties
        h_pbs = 1000;          % [W/m^2K] Convection coefficient of PBS. Measured by aligning simulation
to actual. 590 W/m^2K calculated from Supplemental Figure 2.
```

```
        T_inf_pbs = 37;        % [degC] Environmental temperature

    % Air Properties
        h_air = 23.2;           % [W/m^2K] Convection coefficient of air. Measured by aligning simulation
to actual, Supplemental Figure 1.
        T_inf_air = 37;        % [degC] Environmental temperature

    % Heat Generation Properties
        V = pi*(R^2) * H;      % [m^3] Volume
        f = 1.1;               % [1/s] Loading frequency
        E_v = 142.9*1000;      % [Pa] Viscous loading
        e_0 = .05;             % [-] Maximum compressive strain
        W = (2*pi)*f;          % [(2*pi)/f] Loading period

%% Simulation Set-up
% Simulation Environment
    % Node Properties
        M = 50;             % Nodal divisions in the radial direction
        dr = R/M;           % [m] Spacing between radial nodes

        N = M;              % Nodal divisions in the vertical direction
        dz = H/N;           % [m] Spacing between vertical nodes

    % Simulation Time Properties
        dt = round((dr^2)/(2*alpha),1,'significant');   % [s] Specify the largest possible time step
without causing divergence
        dt = 0.0005; % Manually define dt for testing purposes
        SimTime = 180;   % [s] Cap the total acceptable simulation time
        time = 0;       % [s] Simulation starting time
        MaxSimIter = floor(SimTime/dt);   % Calculate the maximum possible number of iterations
        tarray = linspace(time,SimTime,(SimTime-time)/dt);   % Set the time array for later display

    % Heat Convection Properties
        % These values come from Ozisik Pg 474, and are used for the
        % convection BC's
        beta_pbs = 1 + ((dr * h_pbs)/k);
        gamma_pbs = T_inf_pbs * ((dr * h_pbs)/k);
        r_pbs = (alpha * dt)/(dr^2);

        beta_air = 1 + ((dr * h_air)/k);
        gamma_air = T_inf_air * ((dr * h_air)/k);
        r_air = (alpha * dt)/(dr^2);

    % Heat Generation Properties
        % Calculates Q_dot at every time point
        Q_dot = 4*V*f*E_v*(e_0^2).*sin(W.*tarray).^2;

% Solver and Plotter Properties
    % Solver Properties
        Cont = true;        % Set the "continue" boolean to true
        Temp_Tol = 0.0167;   % Set the temperature significance value [degC/s]
        Tol = abs(T_inf_pbs - T_int)*Temp_Tol;  % Set the value to consider the solution to be SS
        count = 1;          % Allocate the iteration counter

    % Video Properties
        ShowVideo = true;
        framerate = 20; % Set the video playback framerate
        c = floor(1/(framerate*dt)); % Set the video playback quantity

% Simulation Preallocation
    % Create the grid for both the R and H direction for meshing
        [HMat,RMat] = meshgrid(linspace(-R,R,M*2),linspace(0,H,N));

    % Pull out the radius vector for later
        R_Vec = RMat(:,1);

    % Create the initial temperature array and apply the initial condition
    % to it
        T = ones(N,M,1) * T_int;                % Assign a matrix to the initial condition value
```

```
        T_Array = nan(N,M,ceil(SimTime/dt));      % Preallocate the entire temperature array
        T_Array(:,:,count) = T;                 % Assign the first column to the initial temperature
distribution

%% Solver

% Main Iterative Solver Body
while(count < MaxSimIter) && Cont

    % Start off with some housekeeping
        T_old = T;          % Update the new matrix with the old information for this next iteration
        count = count+1;    % Advance the counter. Don't worry, count = 1 is the initial condition, so we
can update it immediately.

    % For the iterative solver, We will work our way inward from the
    % convection boundaries to the zero-flux condition at the center. Let's
    % run the convection condition first. We'll have three different convection boundaries to solve.
(Osizik p475)
        % Upper Air Boundary
        T(1,:) = (2*r_pbs.*T_old(2,:)) + ((1-(2*r_pbs*beta_pbs)).*T_old(1,:)) + (2*r_pbs*gamma_pbs);

        % Lower PBS Boundary
        T(N,:) = (2*r_air.*T_old(N-1,:)) + ((1-(2*r_air*beta_air)).*T_old(N,:)) + (2*r_air*gamma_air);

        % Side PBS Boundary
        T(:,M) = (2*r_pbs.*T_old(:,M-1)) + ((1-(2*r_pbs*beta_pbs)).*T_old(:,M)) + (2*r_pbs*gamma_pbs);

    % Now that the boundary is updated, we can simply step our way into the
    % interior from the outside. Use the old values, of course.
        % Loop over every node from M-1 to node 2
        for i = M-1:-1:2
            for j = N-1:-1:2
            T(j,i) = alpha*dt*((T_old(j,i-1)-2*T_old(j,i)+T_old(j,i+1))/(dr^2)+(T_old(j,i+1)-T_old(j,i-
1))/(2*R_Vec(i)*dr)+((T_old(j-1,i)-2*T_old(j,i)+T_old(j+1,i))/(dz^2))+(Q_dot(count)/k))+T_old(j,i);
            end
        end

    % Finally, update the central point. Since it is a singularity, there
    % is no heat flux. Thus, the temperture is simply the temperature at
    % the adjacent point.
        T(:,1) = T(:,2);

    % Now that iterations are done, lets make a check to see if the
    % temperature change at any point is small enough to consider the
    % system steady-state. There's no need to check simtime, this is
    % included in the while loop.

%          if max(abs(T-T_old))/dt <= Temp_Tol
%              Cont = false;
%          end

    % Now that all of the thinking is done, assign our calculated
    % temperature values to the time-based temperature array
        T_Array(:,:,count) = T;

end

% Once the iterative solver has either converged or time has run out, stop
% the timer
FinalSimTime = toc;

%% Temperature Point-Plot
Line_Plot = figure('Name','Line Simulation','NumberTitle','off');
plot(tarray,squeeze(T_Array(1,1,1:length(tarray))));
hold on
plot(tarray,squeeze(T_Array(round(M/2),1,1:length(tarray))));
plot(tarray,squeeze(T_Array(end,1,1:length(tarray))));
grid on
legend('Bottom Face','Middle Point','Top Face')
```

```
title('Temperature Lines At Boundaries')
xlabel('Time (t) [s]')
ylabel('Temperature (T) [\circC]')
%% Post-Processing
% Check to see if the solution converged. If not, report it.
if ShowVideo
if Cont
    fprintf('Solution has not reached a steady-state value within %is.\n',SimTime)
else
    fprintf('Solution has reached a steady-state value after %fs.\n',tarray(count))
end
% Plotting Setup
    Temperature_Plot = figure('Name','Temperature Simulation','NumberTitle','off');
    set(Temperature_Plot,'WindowStyle','docked');

    tiledlayout(1,2)
    nexttile
    hold on

    axis square
    axis off
    pbaspect([(R*2)/H 1 1])
    set(gca,'nextplot','replacechildren');
    title('Chondrocyte Heat Generation Contour Plot')

    colormap jet
    cbar = colorbar;
    caxis([min([T_int T_inf_pbs T_inf_air]) max([T_int T_inf_pbs T_inf_air])]);
    cbar.Label.String = 'Temperature (T) [\circC]';

    % Access the plot
    pause(0.25)
    figure(Temperature_Plot)


% Contour Plotting
        % To perform the contour plot, we must extend the temperature
        % vector out into a matrix. Perform the plotting at dt speed

            % Cull the NAN's
                T_Array = T_Array(:,:,all(all(~isnan(T_Array))));
                tarray = tarray(~isnan(tarray));

        % Set up the video recorder and plot the results, delayed by the time step
            v = VideoWriter('CHS.mp4','MPEG-4');
            v.FrameRate = framerate;
            open(v);

        for i = 2:c:count
            nexttile(1)
            PlotT = [fliplr(T_Array(:,:,i)) T_Array(:,:,i)];
            surf(HMat,RMat,PlotT,'LineStyle','none');
            view(0,90)

            nexttile(2)
            axis off
            dim = [0.55 0.1 0.4 0.8];

            str = append('---------------Simulation Details---------------',newline,...
                         'Current Time: ',num2str(round(tarray(i),2),'%.2f'),'s',newline,...
                         'Total Simulated Time: ',num2str(tarray(end)),'s',newline,newline,...
                         'Total Iterations: ',num2str(count),newline,...
                         'Total Computation Time: ',num2str(FinalSimTime,2),' s',newline,...
                         'Steady-State: ',mat2str(~Cont),newline,newline,...
                         '--------------Environment Details--------------',newline,...
                         'Material: ',Material,newline,...
                         'C_p: ',num2str(C_p),' kJ/kgK',newline,...
                         'k: ',num2str(k),' W/mK',newline,...
                         'h: ',num2str(h_pbs),' W/m^2K',newline,...
```

```
                    'R: ',num2str(R*1000),' mm',newline,...
                    'H: ',num2str(H*1000),' mm',newline,...
                    'T_i_n_t: ',num2str(T_int),'\circC',newline,...
                    'T_\infty: ',num2str(T_inf_pbs),'\circC',newline...
                    );


            timeann = annotation('textbox',dim,'String',str);
            timeann.FontWeight = 'bold';
            timeann.FontSize = 12;
            timeann.BackgroundColor = [1 1 1];
            timeann.HorizontalAlignment = 'center';
            timeann.VerticalAlignment = 'middle';

            frame = getframe(gcf);
            writeVideo(v,frame);
            delete(timeann);
        end

%% Cleanup

 close(v);   % Close the video writer

 % Clear the figure and prompt the user for playback
    close(Temperature_Plot);    % Clear the figure now that recording is finished

    % Prompt the user for playback
    playback = questdlg('Would you like to display the simulation in real-time?', ...
        'Playback Options','Yes','No','No');

    switch playback
        case 'Yes'
            % Display the video interface and dock it
            video = implay('CHS.mp4',framerate);
        case 'No'

    end
end
```

### MATLAB Convection Coefficient Calculator Code

```
%% Erik Myers
% June Research Lab
% Montana State University

% 2D Transient Heat Conduction Simulation with Heat Generation
% V1.0 10/03/2023

% This code is the third version of the transient heat generation
% simulation, using Osizik's textbook and a modified setup of the
% two-dimensional steady-state heat conduction equation for cylindrical heat
% flow. This particular flavor has been modified to include heat generated
% from viscoelastic loading of the agarose gel.

%% Pre-Prep
clear
clc
close all
warning('off')

tic;    % Start timing the program for display later

%% User Defined Inputs
% This section if for the user to define anything they want about the
% simulation which may be changed frequently.

% Material Properties
    % Agarose Gel Material Properties
        Material = 'Steel';

        switch Material
            case '4.5% Agarose Gel'
            R = (0.25/2) * 0.0254; % [m] Gel radius
            H = (0.5)    * 0.0254; % [m] Gel height
            k = 0.1;               % [W/mK] Thermal conductivity
            C_p = 2850;            % [J/kgK] Specific heat capacity, from the DSC calculations
            rho = 1069;            % [kg/m^3] Density

            a = k/(rho*C_p);   % [m^2/s] Thermal diffusivity

            case 'Aluminum'
            R = (0.562/2) * 0.0254; % [m] Cylinder radius
            H = (2)    * 0.0254;    % [m] Cylinder height
            k = 167;                % [W/mK] Thermal conductivity
            C_p = 896;              % [J/kgK] Specific heat capacity
            rho = 2710;             % [kg/m^3] Density

            a = k/(rho*C_p);   % [m^2/s] Thermal diffusivity

            case 'Steel'
            R = (0.511/2) * 0.0254; % [m] Cylinder radius
            H = (2)    * 0.0254;    % [m] Cylinder height
            k = 51.9;               % [W/mK] Thermal conductivity
            C_p = 486;              % [J/kgK] Specific heat capacity
            rho = 7870;             % [kg/m^3] Density
            a = k/(rho*C_p);    % [m^2/s] Thermal diffusivity

        end

        T_int = 3.3;            % Initial cylinder temperature

    % PBS Material Properties
        h_pbs = 400;           % [W/m^2K] Convection coefficient of PBS. Measured by aligning simulation to actual. 590
calculated.
        T_inf_pbs = 20;        % [degC] Environmental temperature

    % Air Properties
        h_air = h_pbs;           % [W/m^2K] Convection coefficient of air. Measured by aligning simulation to actual.
        T_inf_air = 20;        % [degC] Environmental temperature

    % H value and error vector
        h_low = 10;
        h_high = 50;
        h_mid = (h_high+h_low)/2;
        h_vec = [h_low h_mid h_high];
        SS_Error = [500 1000 1500];
```

```
        ErrorLim = 10;
        Conv_Lim = .01;

    % Heat Generation Properties
        V = pi*(R^2) * H;       % [m^3] Volume
        f = 1.1;                % [1/s] Loading frequency
        E_v = 142.9*1000;       % [kPa] Viscous loading
        e_0 = .05;              % [-] Maximum compressive strain
        W = (2*pi)/f;           % [(2*pi)/f] Loading period

%% Simulation Set-up
% Simulation Environment
    % Node Properties
        M = 50;             % Nodal divisions in the radial direction
        dr = R/M;           % [m] Spacing between radial nodes

        N = M;              % Nodal divisions in the vertical direction
        dz = H/N;           % [m] Spacing between vertical nodes

    % Simulation Time Properties
        dt = round((dr^2)/(2*a),1,'significant');   % [s] Specify the largest possible time step without causing
divergence
        dt = 0.0006;
        SimTime = 30*10;   % [s] Cap the total acceptable simulation time
        time = 0;       % [s] Simulation starting time
        MaxSimIter = floor(SimTime/dt);   % Calculate the maximum possible number of iterations
        tarray = linspace(time,SimTime,(SimTime-time)/dt);   % Set the time array for later display

    % Heat Convection Properties
        % These values come from Ozisik Pg 474, and are used for the
        % convection BC's
        beta_pbs = 1 + ((dr * h_pbs)/k);
        gamma_pbs = T_inf_pbs * ((dr * h_pbs)/k);
        r_pbs = (a * dt)/(dr^2);

        beta_air = 1 + ((dr * h_air)/k);
        gamma_air = T_inf_air * ((dr * h_air)/k);
        r_air = (a * dt)/(dr^2);

    % Heat Generation Properties
        % Calculates Q_dot at every time point
        %Q_dot = 4*V*f*E_v*(e_0^2).*sin(W.*tarray).^2;
        Q_dot = 0 .*tarray;

% Solver and Plotter Properties
    % Solver Properties
        Cont = true;        % Set the "continue" boolean to true
        Temp_Tol = 0.0167;   % Set the temperature significance value [degC/s]
        Tol = abs(T_inf_pbs - T_int)*Temp_Tol;  % Set the value to consider the solution to be SS
        count = 1;          % Allocate the iteration counter

    % Video Properties
        ShowVideo = false;
        framerate = 30; % Set the video playback framerate
        c = floor(1/(framerate*dt)); % Set the video playback quantity

% Data Import and H-value Properties
    Exp_Data = cell2mat(struct2cell(load('Steel_Aircooled.mat')));
    Exp_Data_X = Exp_Data(:,1);
    Exp_Data_Y = Exp_Data(:,2);

    Error_Plot = figure('WindowStyle','docked');
    % Calculate the spline for the exp results
    Exp_XX = round(min(Exp_Data_X)):1:round(max(Exp_Data_X));
    Exp_YY = spline(Exp_Data_X,Exp_Data_Y,Exp_XX);

    % First Plot Boolean
    FP = true;

% Simulation Preallocation
    % Create the grid for both the R and H direction for meshing
        [HMat,RMat] = meshgrid(linspace(-R,R,M*2),linspace(0,H,N));

    % Pull out the radius vector for later
        R_Vec = RMat(:,1);

    % Create the initial temperature array and apply the initial condition
    % to it
        T = ones(N,M,1) * T_int;                % Assign a matrix to the initial condition value
```

```
        T_Array = nan(N,M,ceil(SimTime/dt));    % Preallocate the entire temperature array
        T_Array(:,:,count) = T;                 % Assign the first column to the initial temperature distribution

%% Solver

% Find the error of the low and high H values outside of  te loop, before
% we even start. First, low H error.
count = 1;
T = ones(N,M,1) * T_int;                % Assign a matrix to the initial condition value
T_Array = nan(N,M,ceil(SimTime/dt));    % Preallocate the entire temperature array
T_Array(:,:,count) = T;                 % Assign the first column to the initial temperature distribution

% Calculate new values with the new H
% These values come from Ozisik Pg 474, and are used for the
% convection BC's
beta_pbs = 1 + ((dr * h_vec(1))/k);
gamma_pbs = T_inf_pbs * ((dr * h_vec(1))/k);
r_pbs = (a * dt)/(dr^2);

beta_air = 1 + ((dr * h_vec(1))/k);
gamma_air = T_inf_air * ((dr * h_vec(1))/k);
r_air = (a * dt)/(dr^2);

save('WorkspaceData'); % Save the workspace so the variables can be brought into the function easily

% Run the optimization function
[T,T_Array] = PointOptimization;
save('WorkspaceData'); % Update the saved workspace to have the newly calculated values of T and T_Array in

% Calculate the error at the current H value. First, we create a spline of
%  sim data set to compare them over the same range.
Sim_XX = round(min(tarray)):1:round(max(tarray));
Sim_YY = spline(tarray,squeeze(T_Array(1,1,1:length(tarray))),Sim_XX);

% Now find the bounds for the splines
LB = max([Sim_XX(1) Exp_XX(1)]);
UB = min([Sim_XX(end) Exp_XX(end)]);

% Cull the two data sets based on the index values
Sim_YY = Sim_YY(find(Sim_XX==LB):find(Sim_XX==UB));
Sim_XX = Sim_XX(find(Sim_XX==LB):find(Sim_XX==UB));
Exp_YY = Exp_YY(find(Exp_XX==LB):find(Exp_XX==UB));
Exp_XX = Exp_XX(find(Exp_XX==LB):find(Exp_XX==UB));

% Calculate the actual sos of the error and find the new h value.
SS_Error(1) = sum((Sim_YY-Exp_YY).^2);

% Next, high H error.
count = 1;
T = ones(N,M,1) * T_int;                % Assign a matrix to the initial condition value
T_Array = nan(N,M,ceil(SimTime/dt));    % Preallocate the entire temperature array
T_Array(:,:,count) = T;                 % Assign the first column to the initial temperature distribution

% Calculate new values with the new H
% These values come from Ozisik Pg 474, and are used for the
% convection BC's
beta_pbs = 1 + ((dr * h_vec(3))/k);
gamma_pbs = T_inf_pbs * ((dr * h_vec(3))/k);
r_pbs = (a * dt)/(dr^2);

beta_air = 1 + ((dr * h_vec(3))/k);
gamma_air = T_inf_air * ((dr * h_vec(3))/k);
r_air = (a * dt)/(dr^2);

fprintf('h-value\t|\tSS_Error\t|\tCalculation Time\n');
fprintf('---------------------------------------------\n');
fprintf('%i\t\t\t%0.0f\t\t\t\t%0.0f\n',h_vec(1),SS_Error(1),toc)
tic

[T,T_Array] = PointOptimization; % Run the point optimization function again
save("WorkspaceData.mat");

% Calculate the error at the current H value. First, we create a spline of
%  sim data set to compare them over the same range.
Sim_XX = round(min(tarray)):1:round(max(tarray));
Sim_YY = spline(tarray,squeeze(T_Array(1,1,1:length(tarray))),Sim_XX);

% Now find the bounds for the splines
LB = max([Sim_XX(1) Exp_XX(1)]);
```

```
UB = min([Sim_XX(end) Exp_XX(end)]);

% Cull the two data sets based on the index values
Sim_YY = Sim_YY(find(Sim_XX==LB):find(Sim_XX==UB));
Sim_XX = Sim_XX(find(Sim_XX==LB):find(Sim_XX==UB));
Exp_YY = Exp_YY(find(Exp_XX==LB):find(Exp_XX==UB));
Exp_XX = Exp_XX(find(Exp_XX==LB):find(Exp_XX==UB));

% Calculate the actual sos of the error and find the new h value.
SS_Error(3) = sum((Sim_YY-Exp_YY).^2);

fprintf('%i\t\t%0.0f\t\t\t\t\t%0.0f\n',h_vec(3),SS_Error(3),toc)
%% Main Iterative Solver Body
% Add the while loop for calculating the H value
while SS_Error(2) > ErrorLim && abs(SS_Error(3)-SS_Error(1)) > Conv_Lim
tic
count = 1;
T = ones(N,M,1) * T_int;                % Assign a matrix to the initial condition value
T_Array = nan(N,M,ceil(SimTime/dt));    % Preallocate the entire temperature array
T_Array(:,:,count) = T;                 % Assign the first column to the initial temperature distribution

% Calculate new values with the new H
% These values come from Ozisik Pg 474, and are used for the
% convection BC's
beta_pbs = 1 + ((dr * h_vec(2))/k);
gamma_pbs = T_inf_pbs * ((dr * h_vec(2))/k);
r_pbs = (a * dt)/(dr^2);

beta_air = 1 + ((dr * h_vec(2))/k);
gamma_air = T_inf_air * ((dr * h_vec(2))/k);
r_air = (a * dt)/(dr^2);

[T,T_Array] = PointOptimization;

% Calculate the error at the current H value. First, we create a spline of
%  sim data set to compare them over the same range.
Sim_XX = round(min(tarray)):1:round(max(tarray));
Sim_YY = spline(tarray,squeeze(T_Array(1,1,1:length(tarray))),Sim_XX);

% Now find the bounds for the splines
LB = max([Sim_XX(1) Exp_XX(1)]);
UB = min([Sim_XX(end) Exp_XX(end)]);

% Cull the two data sets based on the index values
Sim_YY = Sim_YY(find(Sim_XX==LB):find(Sim_XX==UB));
Sim_XX = Sim_XX(find(Sim_XX==LB):find(Sim_XX==UB));
Exp_YY = Exp_YY(find(Exp_XX==LB):find(Exp_XX==UB));
Exp_XX = Exp_XX(find(Exp_XX==LB):find(Exp_XX==UB));

% Calculate the actual sos of the error and find the new h value.
SS_Error(2) = sum((Sim_YY-Exp_YY).^2);

if abs(SS_Error(3) - SS_Error(1)) > 10
fprintf('%0.0f\t\t\t%0.0f\t\t\t\t\t%0.0f\n',h_vec(2),SS_Error(2),toc)
else
fprintf('%0.2f\t\t%0.2f\t\t\t\t%0.0f\n',h_vec(2),SS_Error(2),toc)
end
% Calculate new values with the new H
% These values come from Ozisik Pg 474, and are used for the
% convection BC's
beta_pbs = 1 + ((dr * h_pbs)/k);
gamma_pbs = T_inf_pbs * ((dr * h_pbs)/k);
r_pbs = (a * dt)/(dr^2);

beta_air = 1 + ((dr * h_air)/k);
gamma_air = T_inf_air * ((dr * h_air)/k);
r_air = (a * dt)/(dr^2);

% Plot the results. First plot the current H value comparison.
subplot(1,2,1)
plot(Exp_XX,Exp_YY,'r-')
grid on
hold on
title('Simulated/Actual Temperatures')
xlabel('Time (t) [s]')
ylabel('Temperature (T) [\circC]')
plot(Sim_XX,Sim_YY,'b--');
legend('Experimental','Simulation','Location','southeast','AutoUpdate','off')
shg
```

```
hold off

subplot(1,2,2)
if FP
semilogy(h_vec,SS_Error,'k.')
hold on
grid on
FP = false;
end
semilogy(h_vec(2),SS_Error(2),'r.')
semilogy(h_vec(1),SS_Error(1),'k.',h_vec(3),SS_Error(3),'k.')

grid on
hold on
title('Error between Simulated and Actual Temperatures')
xlabel('H-value (h) [W/m^2]')
ylabel('Sum of Squares of Temperature Error (e_S_S) [\circC^2]')
drawnow


% Now pick which endpoint to replace and overwright the two matrices.
% Calculate the slope between the two ends.
E_Slope = [abs((SS_Error(2)-SS_Error(1))/(h_vec(2)-h_vec(1))) abs((SS_Error(3)-SS_Error(2))/(h_vec(3)-h_vec(2)))];

if E_Slope(1) > E_Slope(2)
    h_vec(1) = h_vec(2);
    h_vec(2) = (h_vec(3)+h_vec(1))/2;
    SS_Error(1) = SS_Error(2);
else
    h_vec(3) = h_vec(2);
    h_vec(2) = (h_vec(3)+h_vec(1))/2;
    SS_Error(3) = SS_Error(2);
end

end

FinalSimTime = toc;

fprintf('\nFinal calculated h-value: %0.1f W/m^2.\n',h_vec(2))
%% Post-Processing
% This section was shamelessly stolen directly from V1.0. Don't be upset.
% Check to see if the solution converged. If not, report it.
if ShowVideo
if Cont
    fprintf('Solution has not reached a steady-state value within %is.\n',SimTime)
else
    fprintf('Solution has reached a steady-state value after %fs.\n',tarray(count))
end
% Plotting Setup
    Temperature_Plot = figure('Name','Temperature Simulation','NumberTitle','off');
    set(Temperature_Plot,'WindowStyle','docked');

    tiledlayout(1,2)
    nexttile
    hold on

    axis square
    axis off
    pbaspect([(R*2)/H 1 1])
    set(gca,'nextplot','replacechildren');
    title('Chondrocyte Heat Generation Contour Plot')

    colormap jet
    cbar = colorbar;
    caxis([min([T_int T_inf_pbs T_inf_air]) max([T_int T_inf_pbs T_inf_air])]);
    cbar.Label.String = 'Temperature (T) [\circC]';

    % Access the plot
    pause(0.25)
    figure(Temperature_Plot)


% Contour Plotting
        % To perform the contour plot, we must extend the temperature
        % vector out into a matrix. Perform the plotting at dt speed

            % Cull the NAN's
                T_Array = T_Array(:,:,all(all(~isnan(T_Array))));
                tarray = tarray(~isnan(tarray));
```

```
        % Set up the video recorder and plot the results, delayed by the time step
            v = VideoWriter('CHS.mp4','MPEG-4');
            v.FrameRate = framerate;
            open(v);

        for i = 2:c:count
            nexttile(1)
            PlotT = [fliplr(T_Array(:,:,i)) T_Array(:,:,i)];
            surf(HMat,RMat,PlotT,'LineStyle','none');
            view(0,90)

            nexttile(2)
            axis off
            dim = [0.55 0.1 0.4 0.8];

            str = append('---------------Simulation Details---------------',newline,...
                         'Current Time: ',num2str(round(tarray(i),2),'%.2f'),'s',newline,...
                         'Total Simulated Time: ',num2str(tarray(end)),'s',newline,newline,...
                         'Total Iterations: ',num2str(count),newline,...
                         'Total Computation Time: ',num2str(FinalSimTime,2),' s',newline,...
                         'Steady-State: ',mat2str(~Cont),newline,newline,...
                         '--------------Environment Details--------------',newline,...
                         'Material: ',Material,newline,...
                         'C_p: ',num2str(C_p),' kJ/kgK',newline,...
                         'k: ',num2str(k),' W/mK',newline,...
                         'h: ',num2str(h_pbs),' W/m^2K',newline,...
                         'R: ',num2str(R*1000),' mm',newline,...
                         'H: ',num2str(H*1000),' mm',newline,...
                         'T_i_n_t: ',num2str(T_int),'\circC',newline,...
                         'T_\infty: ',num2str(T_inf_pbs),'\circC',newline...
                         );


            timeann = annotation('textbox',dim,'String',str);
            timeann.FontWeight = 'bold';
            timeann.FontSize = 12;
            timeann.BackgroundColor = [1 1 1];
            timeann.HorizontalAlignment = 'center';
            timeann.VerticalAlignment = 'middle';

            frame = getframe(gcf);
            writeVideo(v,frame);
            delete(timeann);
        end

%% Cleanup

 close(v);   % Close the video writer

 % Clear the figure and prompt the user for playback
    close(Temperature_Plot);    % Clear the figure now that recording is finished

    % Prompt the user for playback
    playback = questdlg('Would you like to display the simulation in real-time?', ...
         'Playback Options','Yes','No','No');

    switch playback
        case 'Yes'
            % Display the video interface and dock it
            video = implay('CHS.mp4',framerate);
        case 'No'

    end
end
```

#### MATLAB Convection Coefficient Calculation Point Optimization Function

```
function [T,T_Array] = PointOptimization
load("WorkspaceData.mat")
    while(count < MaxSimIter) && Cont

        % Start off with some housekeeping
            T_old = T;          % Update the new matrix with the old information for this next iteration
            count = count+1;    % Advance the counter. Don't worry, count = 1 is the initial condition, so we can update
it immediately.
```

```
        % For the iterative solver, We will work our way inward from the
        % convection boundaries to the zero-flux condition at the center. Let's
        % run the convection condition first. We'll have three different convection boundaries to solve. (Osizik p475)
            % Lower PBS Boundary
            T(1,:) = (2*r_pbs.*T_old(2,:)) + ((1-(2*r_pbs*beta_pbs)).*T_old(1,:)) + (2*r_pbs*gamma_pbs);

            % Upper Air Boundary
            T(N,:) = (2*r_air.*T_old(N-1,:)) + ((1-(2*r_air*beta_air)).*T_old(N,:)) + (2*r_air*gamma_air);

            % Side PBS Boundary
            T(:,M) = (2*r_pbs.*T_old(:,M-1)) + ((1-(2*r_pbs*beta_pbs)).*T_old(:,M)) + (2*r_pbs*gamma_pbs);

        % Now that the boundary is updated, we can simply step out way into the
        % interior from the outside. Use the old values, of course.
            % Loop over every node from M-1 to node 2
            for i = M-1:-1:2
                for j = N-1:-1:2
                T(j,i) = a*dt*((T_old(j,i-1)-2*T_old(j,i)+T_old(j,i+1))/(dr^2)+(T_old(j,i+1)-T_old(j,i-
1))/(2*R_Vec(i)*dr)+((T_old(j-1,i)-2*T_old(j,i)+T_old(j+1,i))/(dz^2))+(Q_dot(count)/k))+T_old(j,i);
                end
            end

        % Finally, update the central point. Since it is a singularity, there
        % is no heat flux. Thus, the temperture is simply the temperature at
        % the adjacent point.
            T(:,1) = T(:,2);

        % Now that iterations are done, lets make a check to see if the
        % temperature change at any point is small enough to consider the
        % system steady-state. There's no need to check simtime, this is
        % included in the while loop.

    %         if max(abs(T-T_old))/dt <= Temp_Tol
    %             Cont = false;
    %         end

        % Now that all of the thinking is done, assign our calculated
        % temperature values to the time-based temperature array
            T_Array(:,:,count) = T;

    end
end
```
